# Developmental expression of risk genes implicates the age of onset for neuropsychiatric disorders

**DOI:** 10.1101/2025.08.27.672343

**Authors:** Chao Xue, Xuegao Yu, Lihang Ye, Ying Luo, Liubin Zhang, Bin Tang, Hongsheng Gui, Lin Jiang, Miaoxin Li

**Author notes:** To whom correspondence should be addressed. (L. J.); (M. L.). The authors wish it to be known that, in their opinion, the first two authors should be regarded as joint First Authors.

## Abstract

The functional effects of genetic variants associated with complex diseases exhibit pronounced spatiotemporal specificity. Although spatially resolved studies have advanced, their temporal dynamics remain poorly characterized. Here, we present an analytical framework integrating developmental gene expression with genome-wide association studies to decipher age-specific windows during which genetic variants exert their effects and to elucidate underlying mechanisms. Applying this framework to five major neuropsychiatric disorders, we uncover a fundamental principle: the peak incidence of a disease precisely coincides with the developmental window of peak expression of its associated risk genes in the prefrontal cortex. These risk windows are characterized by distinct biological processes; for instance, childhood risk for attention-deficit/hyperactivity disorder aligns with a peak in presynaptic machinery gene expression, whereas late-life risk for Alzheimer’s disease corresponds to heightened immune-related gene activity. Leveraging this principle of temporal convergence significantly improves the prioritization of disease genes. Our work establishes the developmental basis for the age of onset of complex diseases, providing a temporal roadmap for understanding disease mechanisms and developing age-appropriate therapeutic strategies.

## Introduction

The effects of genetic variants are governed by a fundamental spatiotemporal context, manifesting only when their target genes are expressed at the right time and in the right place^1–3^. While immense progress has been made in mapping the spatial context of disease risk from tissues to single cells^4–8^, the temporal dimension remains poorly resolved. Understanding how the pathogenic effects of genetic risk factors are deployed across the human lifespan is a critical unanswered question in human genetics.

The importance of this temporal dimension is particularly striking in neuropsychiatric disorders, which are characterized by distinct, age-specific windows of onset^9^. Elucidating the mechanisms that determine this age-dependent susceptibility is not only crucial for understanding pathogenesis but also for developing timely interventions that can improve outcomes^10,11^. Yet, systematically mapping this temporal landscape has been intractable. This endeavor has been hindered by the historical scarcity of high-resolution transcriptomic data spanning human development and, critically, by the lack of analytical frameworks capable of integrating dynamic gene expression patterns with summary-level data from genome-wide association studies (GWAS).

Here, we bridge this gap by developing an analytical framework to systematically resolve the age-specific windows of genetic risk for complex traits. We integrate developmental gene expression dynamics from the human prefrontal cortex with large-scale GWAS for five major neuropsychiatric disorders. Through this approach, we aim to identify the critical developmental epochs of disease vulnerability, characterize the specific biological programs active within them, and establish a new temporal dimension for prioritizing disease genes.

## Results

### A framework to resolve the temporal dynamics of disease risk

We began by addressing the hypothesis that the distinct ages of onset for neuropsychiatric disorders are driven by the temporal dynamics of risk-gene expression. To test this, we developed tDESE (temporal-specific Driver-tissue Estimation by Selective Expression), an analytical framework designed to integrate developmental transcriptomics with GWAS data to pinpoint critical windows of disease risk (Fig. 1). The core challenge in this endeavor is to distill true developmental signals from noisy, post-mortem human brain expression data.

**Figure 1.**
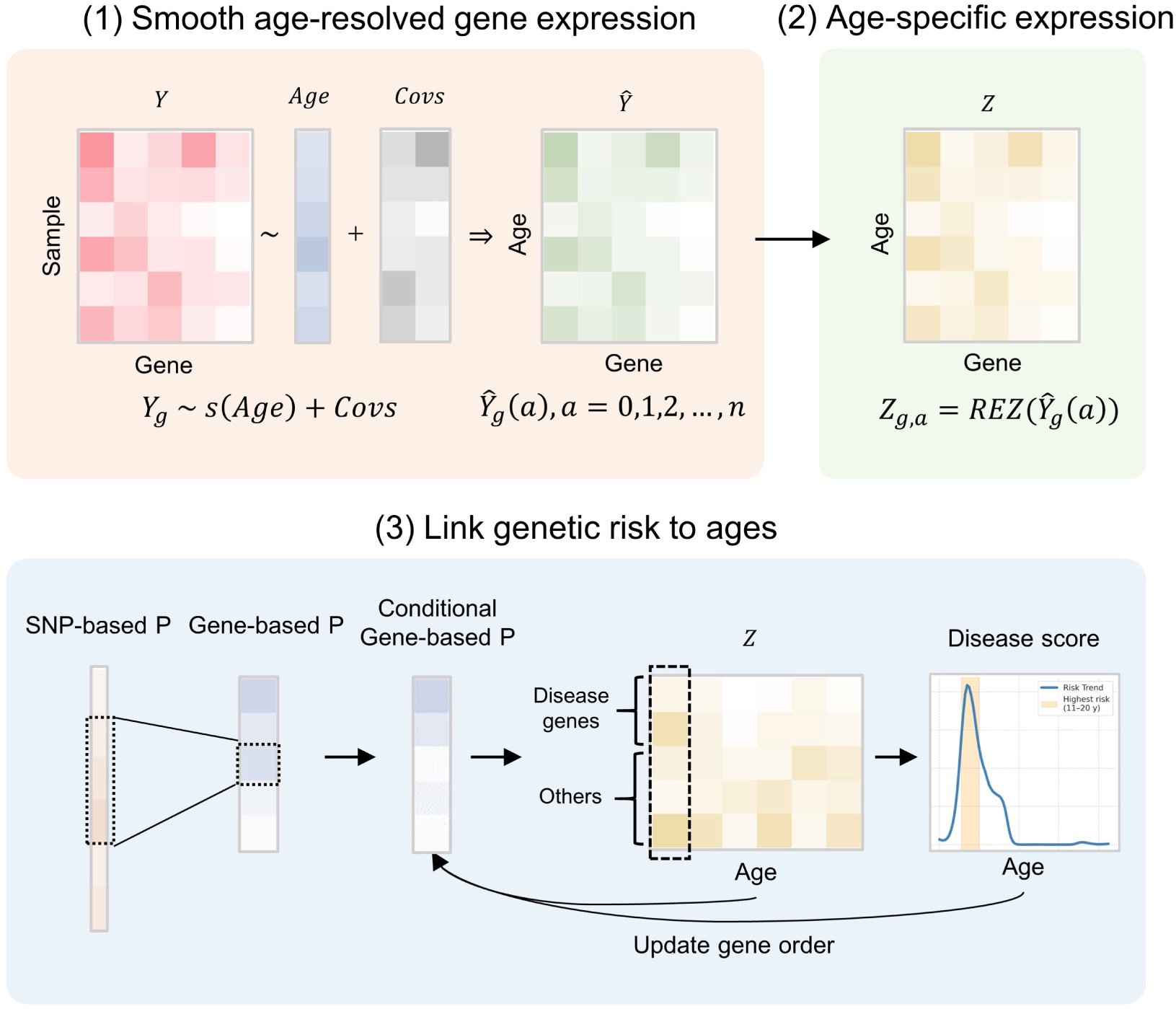
Schematic overview of the tDESE analytical framework. tDESE integrates age-resolved developmental transcriptomic data and disease GWAS summary statistics to identify temporally specific enrichment of genetic risk. The output consists of year-by-year enrichment scores and adjusted P-values for disease association across developmental ages. The framework comprises three major steps: (1) Temporal modeling of gene expression. For each gene, tDESE fits a generalized additive model (GAM) to capture the relationship between expression and age, while adjusting for potential confounders (Covs). Partial dependence analysis is used to estimate the marginal effect of age on expression after controlling for covariates, generating smoothed, high-resolution expression trajectories across age. In the diagram, 𝑌𝑌 denotes the raw expression data, 𝑌𝑌 represents the predicted smoothed expression, 𝑠𝑠(·) denotes the smooth function from GAM, 𝑎𝑎 denotes age, and 𝑔𝑔 denotes gene. (2) Quantification of temporal expression specificity. Relative expression Z-scores are computed to quantify the expression specificity of each gene across developmental time points, yielding an age-specific expression matrix 𝑍𝑍. (3) Enrichment analysis via DESE framework. The age-specific expression matrix 𝑍𝑍 and GWAS P-values are jointly analyzed. SNP-level associations are first aggregated to the gene level. DESE then performs conditional enrichment analysis to test whether genes associated with disease show significantly higher temporal specificity than non-associated genes at each age. An iterative prioritization procedure is applied: in each iteration, genes with strong expression specificity at age windows showing high association are given higher weights, and the process is repeated until convergence. To account for potential biases introduced by the iterative scheme, 100 permutations of the expression matrix are performed to construct a null distribution of enrichment scores, from which corrected P-values are derived. See Methods for full details.

Our first step was therefore to construct robust, high-resolution expression trajectories for each gene across the human lifespan. We proposed a generalized additive model (GAM) to two independent prefrontal cortex developmental datasets, a method chosen for its flexibility in capturing complex, non-linear relationships. This approach allowed us to model gene expression as a continuous function of age while statistically controlling for key technical and biological confounders, such as RNA quality and sex. The result was a set of smoothed expression profiles that represent the dynamic activity of thousands of genes.

With these trajectories established, we next sought to quantify the temporal specificity of each gene’s expression, aiming to identify the precise window of its peak activity. We computed a relative expression Z-score (REZ) for each gene at yearly intervals, which effectively transforms its expression curve into a profile of its functional importance over time. The final and central component of tDESE involves integrating these temporal specificity profiles with

GWAS summary statistics for a complex disease. Extending our previous DESE method^5^, tDESE employs a powerful iterative procedure that refines the set of putative disease-associated genes, substantially enhancing the signal-to-noise ratio. This analysis yields a statistical enrichment score for each developmental time point, pinpointing specific ages where the expression of disease-associated genes is most significantly concentrated, thereby revealing the putative temporal origins of disease pathology.

### Simulations confirm the accuracy and specificity of tDESE

To rigorously validate the statistical performance of tDESE, we conducted extensive simulations using real genotype data from the UK Biobank^12^. We first challenged the framework under a null model, in which the expression of susceptibility genes had no temporal component. As required for a robust method, tDESE produced no spurious signals of enrichment. Across 100 simulations, the mean enrichment scores fluctuated at a low baseline, rarely exceeding a value of 1.5 and showing no discernible trend across the lifespan (Fig. 2a). While a small fraction of individual simulations (4%) yielded a stochastically significant result (adjusted *P* <0.05) at isolated time points, these hits failed to coalesce into any coherent temporal pattern, confirming that the framework’s type I error rate is well-controlled and that it is robust against false positives arising from noise.

**Figure 2.**
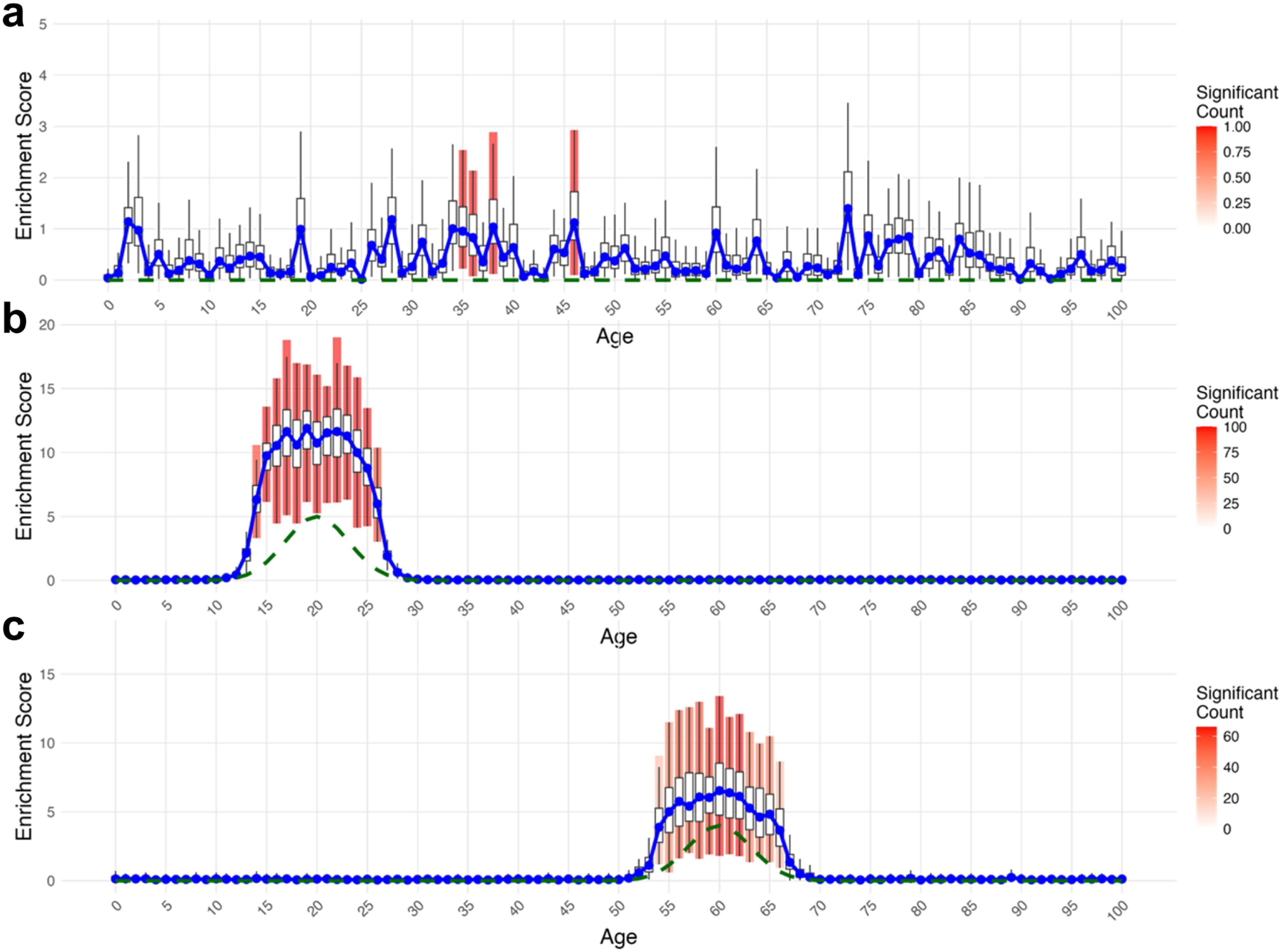
tDESE accurately identifies temporal risk windows in simulations. (a) Under a null model with no temporal gene expression patterns, tDESE shows no spurious enrichment. (b) For a simulated binary trait driven by genes peaking at age 20, tDESE identifies the correct risk window with high power and accuracy. (c) For a simulated continuous trait with risk driven by genes peaking at age 60, the framework again correctly prioritizes the peak age. In all panels, bars represent enrichment scores (red indicates significance at *P* < 0.05), the solid blue line is the mean enrichment score across 100 simulations, and the dashed green line is the simulated mean expression profile of driver genes.

Having established this high specificity, we next tested tDESE’s power to resolve a defined temporal signal. We simulated a binary disease trait (3,000 cases, 3,000 controls) where genetic risk was driven by 250 genes (contributing 25% heritability) whose expression peaked sharply at age 20. The framework resolved this simulated risk window with remarkable precision and power. The resulting profile of mean enrichment scores formed a sharp, high-amplitude peak that precisely mirrored the underlying expression trajectory of the driver genes, soaring from a baseline near zero to a mean score greater than 12 at the epicenter of the risk window (Fig. 2b). This signal was both strong and temporally localized. All 100 simulations returned a highly significant result (adjusted *P* <0.05), with significant enrichment being tightly confined to the peak and its immediately adjacent years (age 20 ± 3 years), a pattern consistent with the high temporal correlation of gene expression.

Finally, to assess the framework’s versatility, we applied it to a more complex and statistically challenging continuous trait within a cohort of 6,000 individuals, with risk driven by an expression peak of 250 genes (contributing 25% heritability) at age 60. Even in this scenario, tDESE successfully identified the simulated risk window. It generated a clear and statistically significant enrichment profile accurately centered on age 60, which displayed the highest average enrichment score (mean = 7.28) (Fig. 2c). As expected for a quantitative trait, the resulting enrichment profile was broader and of lower amplitude than in the binary case, and the statistical power was more modest (66%), yet the signal remained unambiguous. Substantial enrichment was consistently detected in the immediately adjacent ages (58–62), while time points outside the 50–70 year window showed none. Collectively, these simulations establish that tDESE robustly and accurately links the genetic architecture of both categorical and continuous traits to specific developmental epochs.

### Temporal windows of genetic risk for neuropsychiatric disorders align with clinical onset

Having validated tDESE, we applied it to uncover the temporal origins of genetic risk for five major neuropsychiatric disorders with well-characterized onset profiles: attention-deficit/hyperactivity disorder (ADHD)^13^, schizophrenia (SCZ)^14^, major depressive disorder (MDD)^15^, bipolar disorder (BIP)^16^, and Alzheimer’s disease (AD)^17^. These were selected for their pronounced age-specific incidence^9,18^ and the availability of large-scale GWAS (N > 225K per trait). The analysis was conducted using two independent developmental transcriptomic datasets from the dorsolateral prefrontal cortex (DLPFC), referred to as LIBD_szControl and CMC assembled in the PsychENCODE project^19^ (see Methods for details). The prefrontal cortex plays a critical role in higher-order cognitive functions and undergoes prolonged maturation during adolescence and adulthood, a period highly relevant to the etiology of various psychiatric conditions^19^. The datasets offered complementary age coverage, with LIBD_szControl comprising 253 samples from birth to age 71, and CMC comprising 283 samples from age 17 to 90 (Supplementary Fig. 1); and all samples were non-neuropsychiatric controls.

A foundational step was to resolve true developmental signals from the high inter-sample variability inherent in post-mortem tissue. We performed UMAP visualization and pairwise sample correlation analysis on the raw expression data. Samples with similar chronological ages did not exhibit clear expression similarity, indicating high inter-sample variability (Fig. 3a,c; Supplementary Fig. 2a,c), likely due to tissue heterogeneity and inter-individual variation masking age-dependent trends. In contrast, smoothed expression trajectories generated by GAM fitting displayed clear age-continuous clustering (Fig. 3b,d; Supplementary Fig. 2b,d), validating tDESE’s ability to isolate marginal age effects on gene expression.

**Figure 3.**
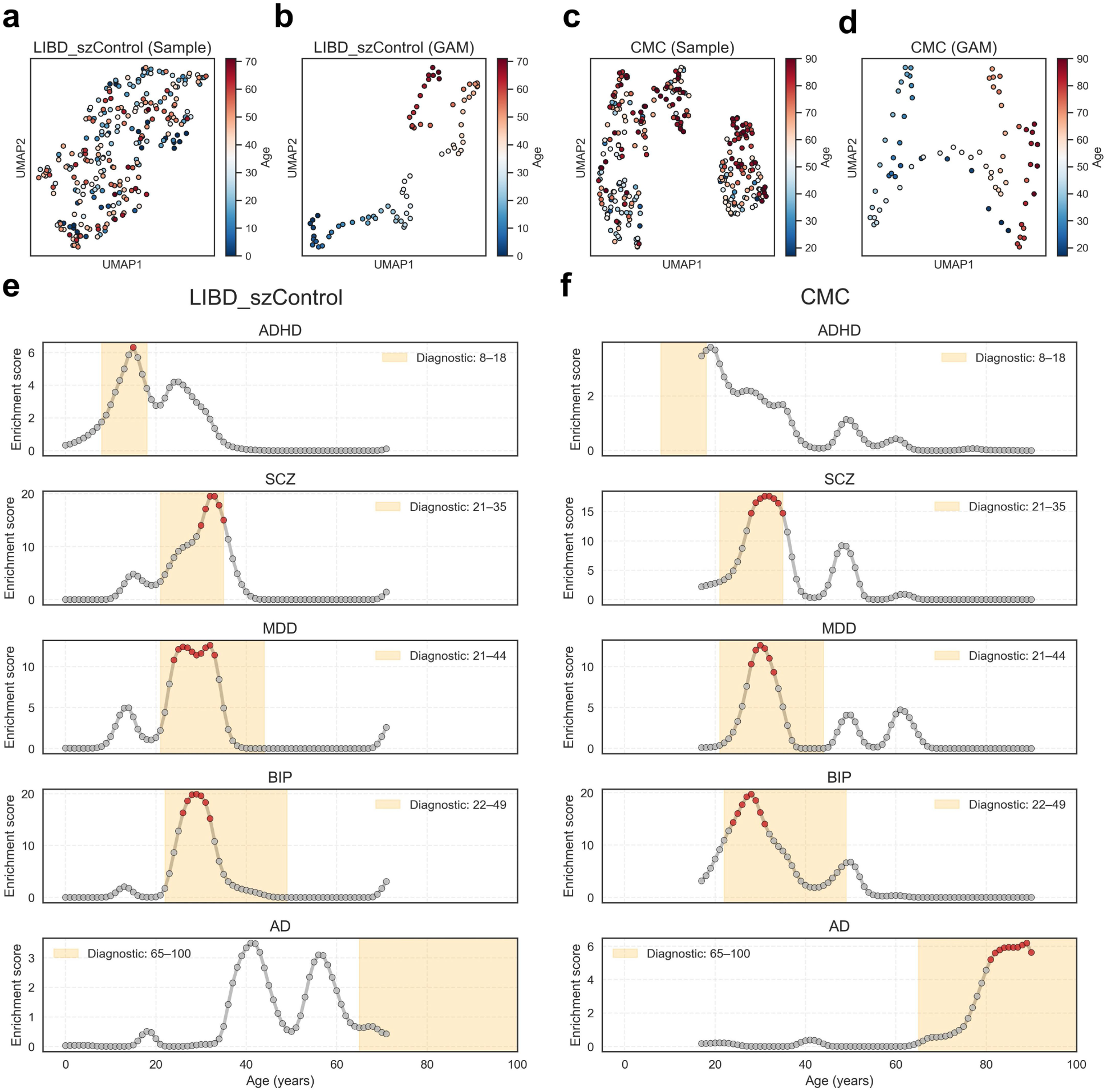
**Estimated age-specific associations for five neuropsychiatric disorders**. tDESE was applied to identify temporally enriched genetic risk windows for five neuropsychiatric disorders with known age-of-onset patterns: ADHD, SCZ, BIP, MDD, and AD. The analysis was performed using two independent developmental gene expression datasets from the dorsolateral prefrontal cortex (DLPFC), namely LIBD_szControl and CMC. After quality control, the LIBD_szControl dataset covered ages 0–71 years, and the CMC dataset spanned ages 17–90 years. (a, c) UMAP visualization based on raw sample-level gene expression profiles. Each point represents a sample, colored by chronological age; axes represent reduced dimensions. (b, d) UMAP visualization based on smoothed, year-specific expression profiles generated via GAM. Each point represents a single year of age, colored by age. (e, f) tDESE-inferred enrichment profiles across age for each of the five disorders. The x-axis indicates age (in years), and the y-axis shows the disease enrichment score. Red dots denote ages with statistically significant enrichment (adjusted *P* < 0.05); grey dots indicate non-significant associations. Yellow shaded regions indicate the reported age- of-onset ranges based on prior literature (see Methods for details).

This approach revealed a striking concordance between the windows of peak genetic risk enrichment and the known clinical age-of-onset for each disorder. For ADHD, a classic childhood-onset disorder, the LIBD_szControl dataset identified a significant enrichment peak for genetic risk at age 15 (adjusted *P* = 0.04), falling squarely within the reported diagnostic window of 8–18 years^9^ (Fig. 3e). In the CMC dataset, no significant ADHD-associated age was detected (Fig. 3f), likely due to the absence of samples younger than 17 years, thereby limiting the detection of early developmental signals.

In contrast, disorders emerging in early to mid-adulthood showed highly concordant risk windows across both independent datasets. For SCZ, tDESE identified a significant risk window spanning ages 30–35 in LIBD_szControl (peak enrichment score = 19.5, adjusted *P* < 0.01) and 28–35 in CMC (peak enrichment score = 17.6, adjusted *P* < 0.01), both aligning precisely with its established onset period of 21–35 years^9^ (Fig. 3e,f). A similar high degree of consistency was observed for MDD, with significant windows at 24–33 years in LIBD_szControl and 28–33 years in CMC, and for BIP, with windows at 26–32 years in LIBD_szControl and 26–31 years in CMC. For both disorders, the peak enrichments were highly significant (adjusted *P* < 0.01) and consistent with their respective clinical onset ranges^9^.

Finally, we examined Alzheimer’s disease, a canonical late-onset neurodegenerative disorder. The results again underscored the framework’s temporal specificity. No signal was detected in the LIBD_szControl cohort, whose age range does not extend into the primary AD risk period. However, in the CMC cohort, tDESE identified a clear, late-life risk window, with enrichment scores escalating sharply after age 80 and peaking at age 89 (adjusted *P* < 0.01) (Fig. 3f). This finding directly mirrors epidemiological data, such as a meta-analysis reporting a rise in AD prevalence in China from 1.3% at ages 65–69 to 18.5% at ages 85–89^20^.

Taken together, our analysis establishes a direct and robust link between the developmental timing of risk-gene expression and the age-specific clinical manifestation of five distinct neuropsychiatric disorders. The findings demonstrate that genetic risk is not a static property but a dynamic process, one that is activated within specific temporal windows to shape the brain’s vulnerability to disease.

### Disease risk windows reveal distinct biological programs

Having identified these disease-specific risk windows, we sought to define the biological processes active within them. We developed a temporal correlation analysis to identify "temporally sensitive genes," referring to genes whose expression trajectories are highly correlated with a disorder’s tDESE enrichment profile (see Methods). Functional analysis of these genes revealed that each risk window is characterized by a distinct and functionally coherent biological program.

Gene Ontology (GO) enrichment analysis of the top 200 temporally sensitive genes for each disorder revealed a fundamental divergence between the psychiatric and neurodegenerative conditions (Fig. 4a). For ADHD, SCZ, MDD, and BIP, the most significantly enriched Cellular Component term was uniformly synapse, establishing synaptic biology as a shared cornerstone of their temporal risk^21–23^. In stark contrast, the late-life risk window for Alzheimer’s disease was dominated by genes involved in the inflammatory response (Biological Process, adjusted *P* = 2.6 × 10⁻⁹), consistent with the well-established role of neuroinflammation in its pathogenesis^24^.

**Figure 4.**
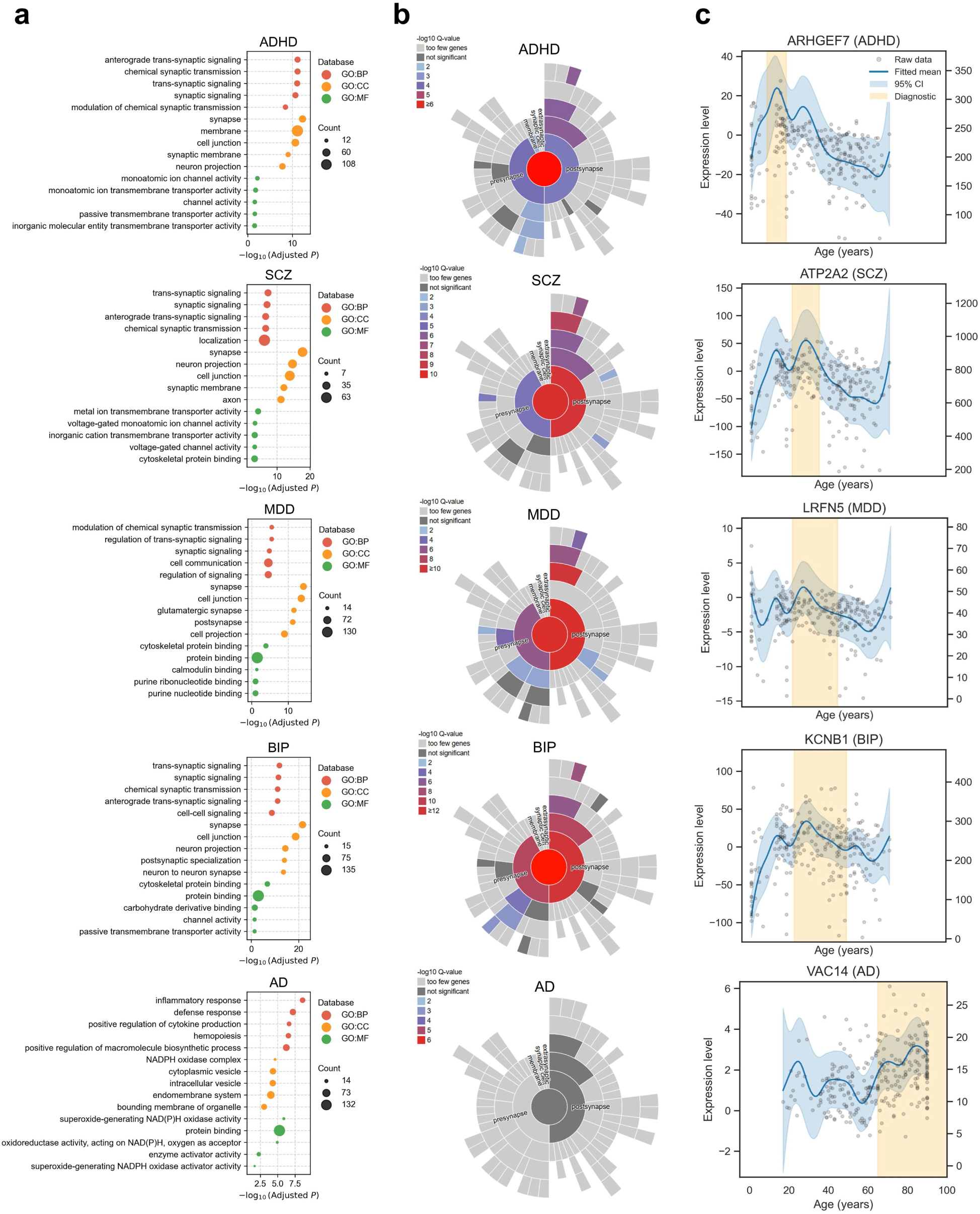
Functional and molecular signatures of disease-specific risk windows. (a) Gene Ontology (GO) enrichment for the top 200 genes temporally correlated with each disorder’s risk profile. Dot size is proportional to the number of genes in a term; color denotes the GO domain (BP, Biological Process; CC, Cellular Component; MF, Molecular Function). (b) Synaptic Gene Ontology (SynGO) analysis reveals compartment-specific enrichment. Color intensity in the circular plot corresponds to statistical significance (-log₁₀ adjusted P-value). (c) Expression trajectories of representative, high-confidence risk genes whose expression dynamics correlate with disease risk. Grey points are raw expression values from individual samples (scaled to the right y-axis); the blue curve is the GAM-smoothed mean expression trajectory (95% CI shaded; scaled to the left y-axis). The yellow shaded area denotes the clinical onset window. Analyses for ADHD, SCZ, MDD, and BIP used the LIBD_szControl dataset; AD analysis used the CMC dataset due to the absence of post-71-year samples in LIBD_szControl.

Within the overarching synaptopathy of the psychiatric disorders, the specific biological processes were distinct. ADHD’s early-life window was most significantly enriched for anterograde trans-synaptic signaling (adjusted *P* = 6.8 × 10⁻¹²). The adult-onset disorders also centered on synaptic transmission, but with different emphases: trans-synaptic signaling was the top term for both SCZ (adjusted *P* = 5.3 × 10⁻⁸) and BIP (adjusted *P* = 1.3 × 10⁻¹²), while MDD was defined by modulation of chemical synaptic transmission (adjusted *P* = 3.3 × 10⁻⁶), reinforcing the synaptopathy model across these conditions^21,23^.

To resolve these synaptic signatures with higher spatial precision, we used the SynGO database^25^, revealing a clear divergence in the implicated synaptic compartments (Fig. 4b). The ADHD risk window was characterized by temporally sensitive genes enriched in both presynaptic (adjusted *P* = 1.1 × 10⁻⁴) and postsynaptic (adjusted *P* = 2.3 × 10⁻⁴) components. In contrast, the risk windows for SCZ, MDD, and BIP were all dominated by a much stronger enrichment for postsynaptic genes. For SCZ, the postsynaptic enrichment (adjusted *P* = 3.2 × 10⁻¹⁰) was five orders of magnitude more significant than the presynaptic one (adjusted *P* = 2.2 × 10⁻⁵). This marked postsynaptic skew was also observed for MDD (adjusted *P* = 2.3 × 10⁻¹¹ vs. 9.9 × 10⁻⁷) and BIP (adjusted *P* = 1.2 × 10⁻¹² vs. 3.9 × 10⁻¹⁰). This suggests a pathogenic model where distinct synaptic vulnerabilities are engaged at different developmental stages.

These overarching signatures are exemplified by the expression dynamics of individual, high-confidence risk genes (Fig. 4c). The ADHD risk window aligns with the adolescent peak expression of *ARHGEF7*, which encodes β-PIX, a key regulator of Rac1/Cdc42 GTPases essential for dendritic spine formation^26^. For SCZ, the risk window coincides with the peak expression of *ATP2A2*, which encodes the calcium pump SERCA2, a critical regulator of the intracellular calcium signaling implicated in the disorder^27^. Similarly, the MDD and BIP risk windows align with the peak expression of genes crucial for synaptic plasticity (*LRFN5*)^28^ and neuronal excitability (*KCNB1*, encoding the Kv2.1 channel)^29^. Finally, the late-life risk for AD is marked by peak expression of *VAC14*, part of the PIKfyve complex essential for PI(3,5)P₂ biosynthesis and lysosomal-autophagic function, whose disruption causes neurodegeneration^30^.

In summary, this integrative analysis identifies temporally sensitive genes that orchestrate age-specific biological processes. These findings provide a new temporal dimension for understanding disease, moving from broad synaptopathy to specific compartmental and molecular vulnerabilities that underpin the age-specific manifestation of neuropsychiatric and neurodegenerative disorders.

### Temporal risk profiles are sexually dimorphic

Given the known sex differences in the prevalence and onset of many psychiatric disorders, we investigated whether tDESE could resolve sex-specific temporal risk profiles. We stratified the analysis for SCZ, MDD, and BIP, the disorders with sufficient sample sizes for this finer-grained test. The analysis revealed a consistent pattern of earlier- emerging genetic risk windows in males compared to females across all three disorders (Fig. 5a).

**Figure 5.**
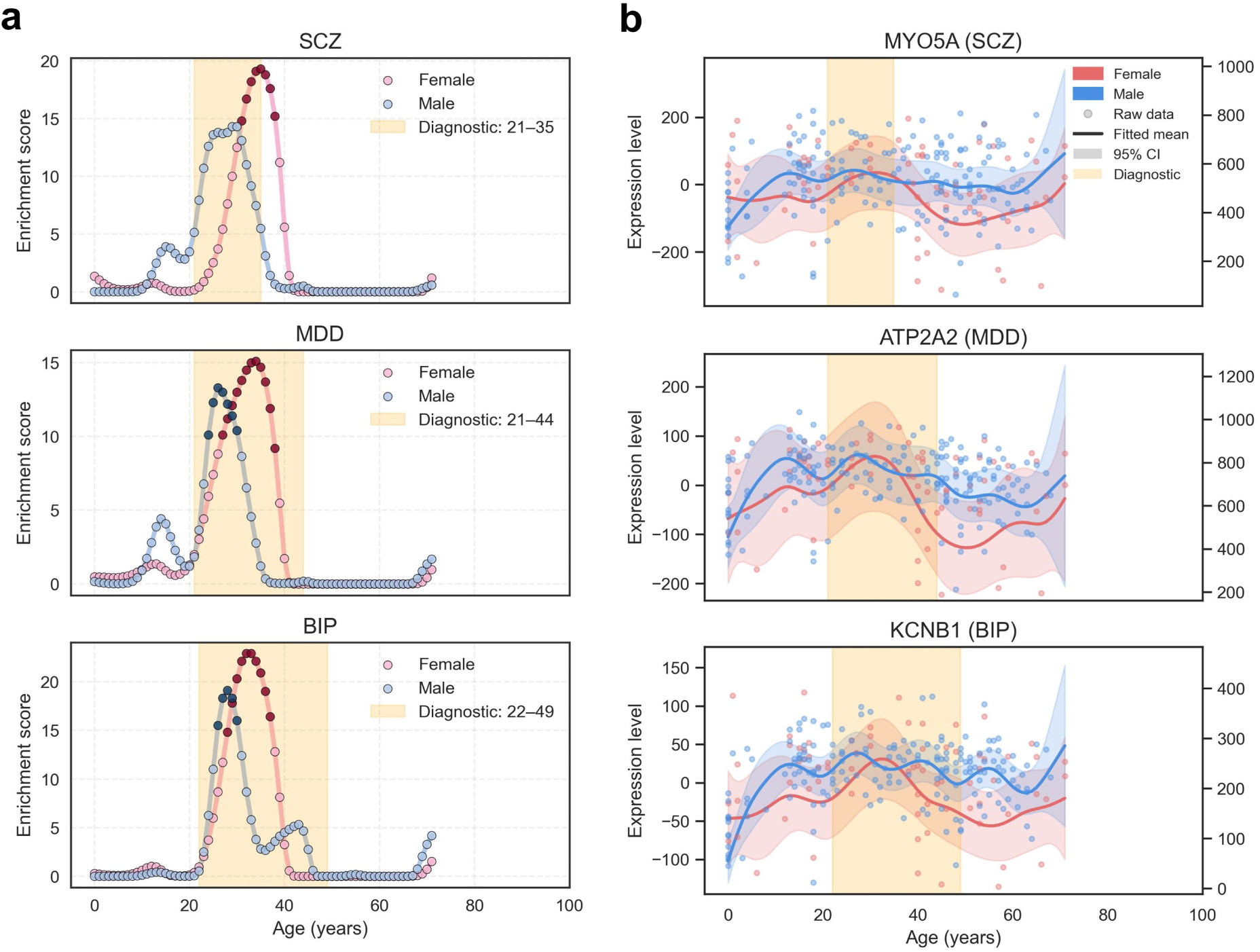
Sexually dimorphic risk windows and gene expression in psychiatric disorders. (a) Sex-stratified tDESE enrichment profiles for SCZ, MDD, and BIP. Female (pink) and male (blue) profiles are shown. Darker points denote statistically significant enrichment (adjusted *P* < 0.05). The shaded yellow area represents the overall clinical onset window. (b) Sex-specific expression trajectories for representative temporally sensitive genes. Curves show GAM-smoothed mean expression for females (red) and males (blue) (95% CI shaded; scaled to the left y-axis). Raw data points for individuals are shown in the background (scaled to the right y-axis). The temporal architecture of genetic risk extends to cognitive and behavioral traits.

For SCZ, we identified a significant risk window for females between ages 31 and 38 (peaking at 35, adjusted *P* < 0.01), whereas males showed an earlier, suggestively significant window at ages 29–30 (adjusted *P* = 0.07). This 3- to-5-year lag in the female risk profile is highly consistent with long-standing epidemiological observations of a later age of onset for SCZ in women^31,32^. A similar temporal shift was observed for BIP, where a significant male risk window from 26 to 30 years (peak at 28, adjusted *P* < 0.01) preceded the female window of 28 to 37 years (peak at 32, adjusted *P* < 0.01). This again aligns with clinical reports of earlier onset in males, particularly for Bipolar I disorder^33^.

For MDD, tDESE also identified an earlier male risk window at 24 to 30 years (peak at 26, adjusted *P* < 0.01), followed by a later and more prolonged female window at 27 to 38 years (peak at 34, adjusted *P* < 0.01). This finding contributes to a complex literature on sex differences in MDD onset^9,34^.

Crucially, these sexually dimorphic risk profiles were mirrored by the expression dynamics of temporally sensitive genes (Fig. 5b). In SCZ, for instance, the expression of *MYO5A* shows an earlier rise and peak in males that coincides with their earlier risk window. Likewise, the expression trajectories of *ATP2A2* in MDD and *KCNB1* in BIP both show earlier up-regulation in males, providing a potential molecular basis for the earlier emergence of genetic risk.

Together, these analyses suggest that sex differences in the clinical manifestation of psychiatric disorders are not merely a clinical phenomenon but may be rooted in sexually dimorphic timing of developmental gene expression programs within the prefrontal cortex. While these findings require validation in larger, sex-stratified cohorts due to the reduced statistical power of this sub-analysis, they demonstrate the remarkable ability of our temporal framework to resolve fine-grained, clinically relevant patterns of disease risk.

### The temporal architecture of genetic risk extends to cognitive and behavioral traits

To determine if the principle of temporally-specific genetic risk extends beyond clinical disorders, we applied tDESE to four major cognitive and behavioral traits: neuroticism (NEU)^35^, smoking behavior (SMK)^36^, sleep duration (SD)^37^, and intelligence (IQ)^38^. This analysis revealed that these traits are also characterized by distinct developmental windows of genetic enrichment, primarily concentrated in adolescence and early adulthood (Fig. 6a).

**Figure 6.**
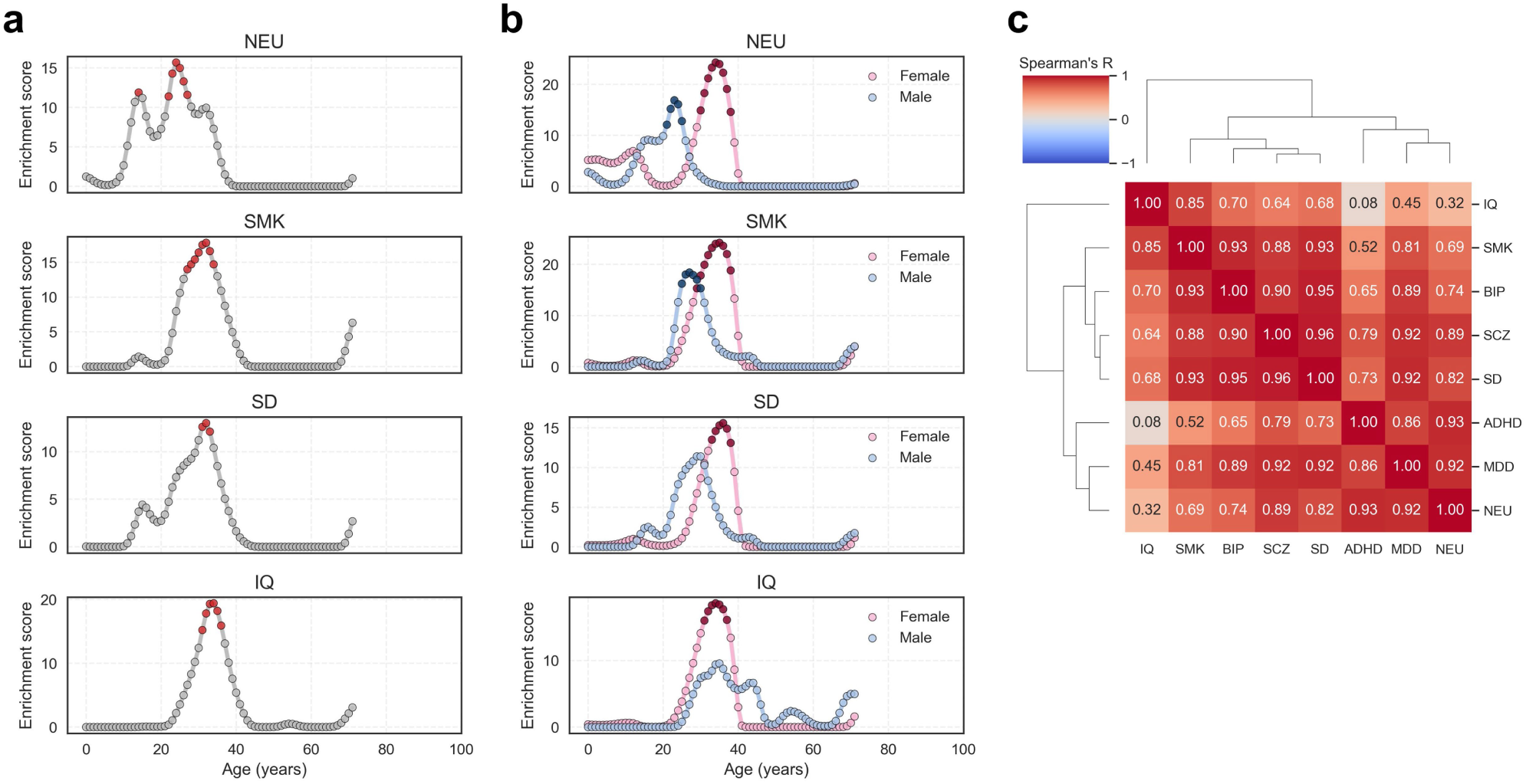
Temporal architecture of genetic risk for cognitive and behavioral traits. (a) Age-specific enrichment profiles for neuroticism (NEU), smoking behavior (SMK), sleep duration (SD), and intelligence (IQ). Red points indicate significant enrichment (adjusted *P* < 0.05). (b) Sex-stratified enrichment profiles for the four traits. Female (pink) and male (blue) profiles are shown; darker points denote statistical significance. (c) Heatmap of Spearman’s correlation between the temporal enrichment profiles of all eight phenotypes. Dendrograms represent hierarchical clustering based on correlation patterns.

Neuroticism was unique in displaying a bimodal risk profile, with a significant adolescent peak at age 14 and a second, larger peak in early adulthood at age 24 (adjusted *P* < 0.01). Sex-stratified analysis revealed a complex and divergent temporal pattern: the adolescent peak appeared earlier in females, while the adult peak emerged significantly earlier in males (age 23 vs. 34 in females, adjusted *P* < 0.01) (Fig. 6b). This developmental "cross-over" suggests that the genetic pathways influencing neuroticism are activated with different sex-specific timing across maturation. In contrast, SMK and SD showed significant enrichment windows in early adulthood (peaks at ages 32 and 32, respectively), and both displayed the earlier male enrichment pattern seen in the psychiatric disorders (Fig. 6b). The genetic architecture of IQ was also temporally localized, with a significant enrichment window at ages 31–36, but showed no significant sex differences.

To evaluate the temporal concordance between cognitive/behavioral traits and psychiatric disorders, we computed Spearman correlations between their genome-wide enrichment curves across all ages (Fig. 6c). NEU displayed high correlation with ADHD (r = 0.93) and MDD (r = 0.92), highlighting shared age-specific genetic architectures^39–41^. SMK was most strongly correlated with BIP (r = 0.93), supporting a shared developmental window for genetic liability to bipolar disorder and addictive behaviors^42^. SD showed strong correlations with SCZ (r = 0.96), BIP (r = 0.95), and MDD (r = 0.92), implicating sleep-regulating pathways in the early developmental origins of multiple psychiatric conditions. In contrast, IQ showed comparatively weaker correlations with all disorders (maximum r = 0.7), suggesting a more distinct genetic architecture.

### Age-specific gene expression enhances the prioritization of known disease genes

We evaluated the performance of age-specific expression profiles for ranking known disease susceptibility genes. Traditional approaches to gene annotation often rely on gene expression data averaged across the brain or developmental stages, potentially masking dynamic molecular signatures associated with specific windows of disease vulnerability. To address this, we used age-specific gene expression levels at each age point to rank candidate genes and evaluated the ability of each age-specific ranking to recover known disease-gene associations. Benchmark gene sets were compiled from the Open Targets Platform^43^, including genes with drug target evidence or high integrative scores from the platform (see Methods for details). The prioritization performance at each age was quantified using the area under the receiver operating characteristic curve (AUC).

For all five neuropsychiatric disorders examined, the highest prioritization performance occurred near the age range of peak disease vulnerability (Fig. 7): ADHD showed the best performance at age 25 (AUC = 0.77), SCZ at age 25 (AUC = 0.61), MDD at age 24 (AUC = 0.67), BIP at age 29 (AUC = 0.61), and AD at age 87 (AUC = 0.62). While the optimal age for ADHD slightly deviated from its epidemiological peak onset, the overall trend suggests that gene expression profiles during high-risk developmental windows more effectively capture pathogenic signals. These findings provide direct support for a key principle: genes with elevated expression during a disorder’s risk period are more likely to be core contributors to disease pathogenesis. This reinforces the notion of an age-dependent genetic architecture in psychiatric disorders and highlights the value of incorporating developmental timing into functional annotation and disease gene discovery. Integrating age-specific expression profiles into future analyses may enhance biological interpretability and improve the translational relevance of gene prioritization frameworks.

**Figure 7.**
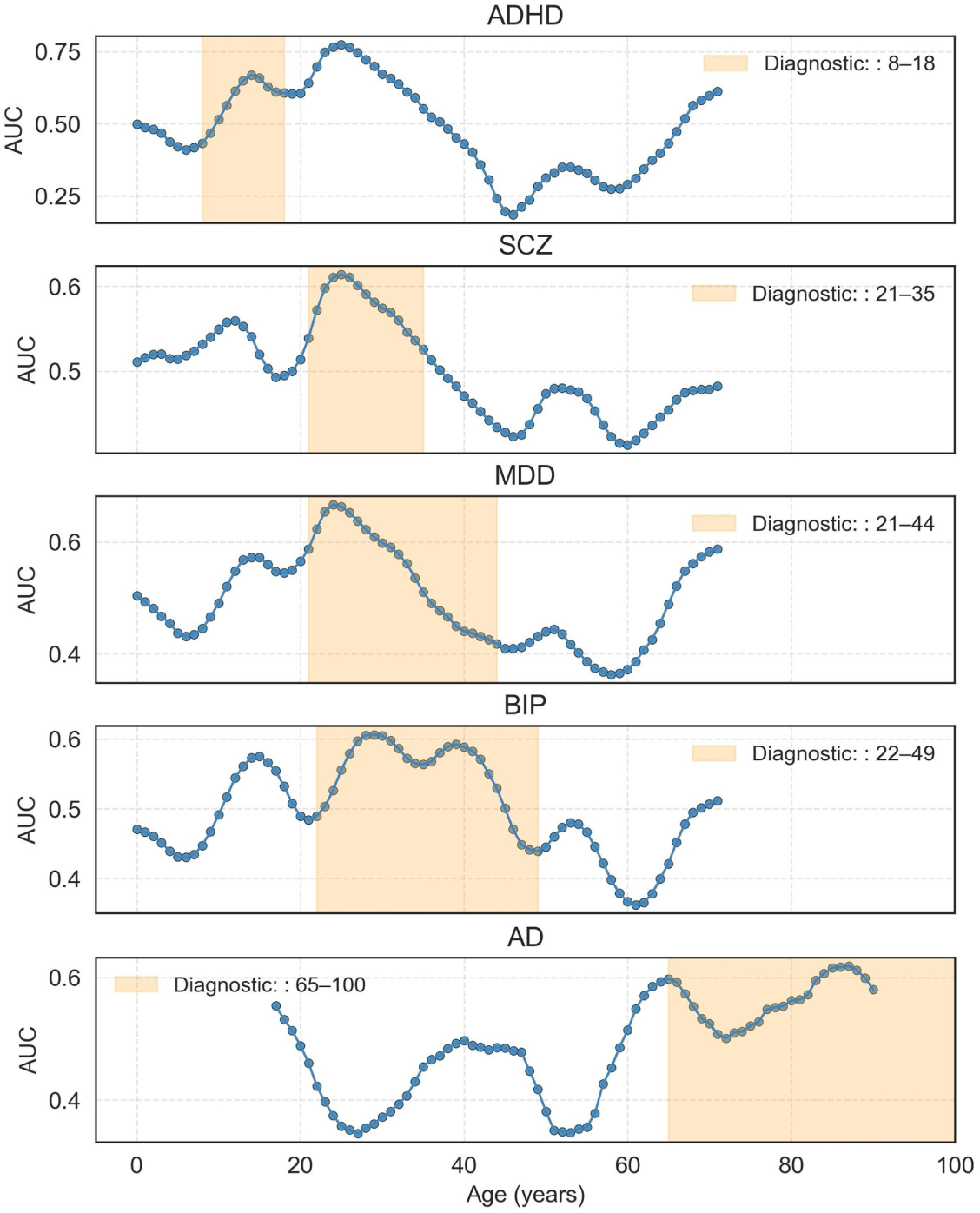
Accuracy evaluation of gene prioritization based on age-specific expression profiles for neuropsychiatric disorders. The ability to prioritize benchmark disease genes (curated from the Open Targets platform) was quantified at each age using the area under the ROC curve (AUC). The y-axis shows AUC performance, plotted against age on the x-axis. For all five disorders, prioritization performance peaks within or adjacent to the clinically defined onset window (shaded). Analyses for ADHD, SCZ, MDD, and BIP used the LIBD_szControl dataset; AD analysis used the CMC dataset.

## Discussion

The timing of disease onset has long been a central mystery in medicine. Here we establish a fundamental principle that helps resolve it: the genetic risk for major neuropsychiatric disorders is activated within specific developmental windows that coincide with the peak expression of associated risk genes. This work moves beyond the prevailing spatially dominated view of genetic risk, recently advanced by spatial transcriptomics^8,44^, to introduce a critical temporal dimension. By integrating developmental transcriptomics with large-scale GWAS, we demonstrate that the vulnerability of a tissue to disease is not static, but is a dynamic property defined by its age-specific functional state. This is exemplified by the alignment of ADHD risk with a period of intense presynaptic maturation and AD risk with an epoch of heightened neuroimmune activity.

Our temporal framework provides a new lens through which to interpret the shared and divergent biology of these disorders. It refines the broad "synaptopathy" model of psychiatric illness^23^ by revealing that different synaptic compartments are implicated at different life stages: a balanced pre- and post-synaptic vulnerability in childhood-onset ADHD gives way to a predominantly postsynaptic vulnerability in adult-onset disorders like schizophrenia. This temporal resolution provides a potential mechanistic basis for their distinct clinical presentations and developmental trajectories. By showing that the same tissue, the prefrontal cortex, can harbor the latent risk for vastly different disorders, our findings underscore that the biological context conferred by age is as critical as the context conferred by anatomy.

A key implication of our work is that the common practice of using static or time-averaged expression data to interpret genetic risk may obscure the most critical pathogenic signals. We provide direct, quantitative evidence for this by showing that the ability to prioritize known causal genes and drug targets is itself temporally specific, peaking precisely within the disease-relevant age window. This provides a data-driven strategy for enriching GWAS follow-up studies and for identifying the most promising targets for age-appropriate therapeutic intervention, addressing a key challenge in post-GWAS analysis^45^.

Our study has limitations that define clear next steps. The temporal transcriptomic datasets, while the best available, are of modest size, and the sex-stratified analyses, though revealing, will require validation in larger cohorts. The observed sexually dimorphic risk profiles, however, are consistent with known differences in brain development driven by the temporal regulation of gene expression by sex hormones^46^. Our analysis is currently focused on the prefrontal cortex; extending this framework to other tissues and a broader range of complex diseases as new longitudinal datasets emerge will be a critical future direction. Finally, while our approach identifies powerful temporal associations, establishing definitive causality is a major challenge in genomics^7,8^. Recognizing these constraints underscores the importance of integrating richer temporal datasets and causal inference strategies to fully realize the potential of time-aware approaches in elucidating complex disease mechanisms.

## Methods

### tDESE framework

#### Overview

The temporal-specific Driver-tissue Estimation by Selective Expression (tDESE) framework integrates temporal gene expression dynamics with GWAS data to identify putative critical developmental windows underlying disease onset and the key genes. By evaluating the age-specific expression patterns of genes linked to disease-associated genetic variants, tDESE pinpoints temporal intervals during which genetic risk may exert maximal influence. Built upon our previously developed DESE framework, tDESE replaces tissue-specific expression profiles with temporally resolved expression data. However, the key challenge is to address inter-sample variability inherent to developmental transcriptomics for a smooth, high-resolution expression trajectory, preferably, at single-year granularity. The variability can be caused by a series of confounding factors, such as uneven sampling density across ages, large inter-sample variability, and non-linear expression changes. Therefore, we developed a generalized additive model (GAM)-based smoothing strategy. This approach enables robust estimation of gene expression trajectories across ages under high-noise conditions, thereby providing a refined signal for identifying developmentally specific genetic risk windows.

The core tDESE workflow consists of three sequential steps: (1) Smoothing age-dependent gene expression trajectories to predict expression levels for each year of age. (2) Calculating the time-specific expression of each gene. (3) Integrating age-specific expression profiles with GWAS data using the DESE framework to identify disease-associated ages.

#### Data input

The inputs to tDESE include GWAS summary statistics, temporal gene expression data, and genotype data from a reference population. For each sample in the temporal expression dataset, the exact age in years is recorded together with relevant covariates, including sex and RNA integrity number (RIN).

#### Expression trajectory modelling and prediction

Because our study focuses on age-dependent dynamic changes in gene expression, we first applied the trimmed mean of M-values (TMM) method to normalize raw expression values across samples, ensuring comparability between samples. As gene expression changes with age are not linear, we used a smoothing approach to model the relationship between gene expression and age. To account for non-age-related sources of variation, sex and RIN were included as covariates in the model. Notably, when modelling was performed separately by sex, sex was naturally excluded as a covariate. tDESE fits a GAM model for each gene individually:

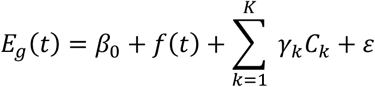

Here, 𝐸𝐸_𝑔𝑔_(𝑡𝑡) denotes the expression level of gene 𝑔𝑔 at age 𝑡𝑡, 𝑓𝑓(𝑡𝑡) is the smoothing function of age implemented using penalized B-splines (P-splines), 𝐶𝐶_𝑘𝑘_ represents the 𝑘𝑘-th covariate including sex and RIN, 𝛾𝛾_𝑘𝑘_ is its regression coefficient, and 𝜀𝜀 is the error term. For categorical covariates, such as sex, values were first converted to dummy variables before linear fitting. The model was implemented using the *pygam* package (version 0.9.1) in Python. The smoothing function parameters followed the default settings in *pygam*, with the number of basis functions (*n_splines*) set to 20 and the penalty strength (*lam*) set to 0.6.

After model fitting, we used partial dependence analysis to extract the marginal effect of age on gene expression while controlling for covariates. We then predicted expression values for each integer age, from the minimum to the maximum sample age, to generate a smoothed expression trajectory at one-year resolution together with the corresponding 95% confidence interval and standard error (SE), which were used to assess the stability and uncertainty of the estimates. This step reduces noise while preserving the authentic signal of temporal dynamic changes as much as possible.

#### Calculation of age-specific expression

To quantify the expression specificity of genes across different ages, we employed the robust-regression z-score (REZ) method^5^. The core concept of this method is to measure the relative expression level of a gene at a given age, defined as its standardized difference compared with other ages. REZ uses robust regression to reduce the influence of outliers, thereby improving the stability of the estimates. In addition, the standard error of expression is incorporated as an indicator of uncertainty, enabling a more reliable representation of the age specificity of gene expression.

#### Linking disease risk to age

We next used the tDESE framework to integrate age-specific gene expression profiles with GWAS summary statistics for each disease, to decipher ages of elevated disease risk. In the original DESE framework, tissue-specific expression profiles are used as input; here, they were replaced by age-specific expression profiles. For a given GWAS dataset, DESE first calculates gene-based association P-values using the ECS statistical model^47^, based on the GWAS P-values of SNPs located within 1 kb upstream or downstream of each gene. We then selected significant genes (Benjamini–Hochberg adjusted *P* < 0.01; if more than 1,000 genes met this criterion, only the top 1,000 by P value were retained) for conditional gene-based association analysis. Specifically, genes were ranked by their gene-based association P values, and conditional gene-based association analysis was performed to remove redundant associations, yielding a set of independently associated genes.

For each age, we used the Wilcoxon rank-sum test to assess whether the age-specific expression of the independently associated genes was higher than that of other genes. The base-10 negative logarithm of the test P-value was defined as the disease enrichment score, which indicates the gene-level association significance of age-specific expression and the disease risk. Based on these enrichment scores and the selective expression profiles of genes, disease-associated genes were re-prioritized, assigning higher priority to genes with high specific expression at disease-enriched ages. Conditional association analysis was then repeated to obtain a new set of disease genes and updated enrichment scores. This procedure was iterated until the results converged.

Because the iterative procedure can introduce potential selection bias for the Wilcoxon rank-sum test, DESE employs a permutation-based global test (maxT permutation test) to estimate the null distribution. Specifically, the age labels in the input age-specific expression profiles are randomly permuted *B* times (100 by default). For each permutation, the analysis is repeated, and the maximum enrichment score (max-ES) across all ages is recorded. These max-ES values form the empirical distribution under the null hypothesis and are used to directly evaluate the significance of the observed statistics. For each age, the permutation P-value is defined as the proportion of values in the null distribution that are greater than or equal to the observed statistic:

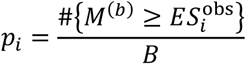

Here, 𝑀𝑀^(𝑏𝑏)^ denotes the maximum enrichment score across all ages in the 𝑏𝑏-th permutation. Because this approach compares the observed statistic with the global maximum from each permutation, the resulting P-values inherently control the family-wise error rate (FWER) and require no additional multiple-testing correction.

In this study, linkage disequilibrium calculations were performed using Phase 3 genotype data from the 1000 Genomes Project, matched to the ancestry of the GWAS samples.

### Simulations

#### Simulation models of gene expression and disease risk

We first modeled the age-dependent expression of a given gene. Its mean expression level, 𝑓𝑓(𝑥𝑥), as a function of age 𝑥𝑥, is described by a Gaussian function:

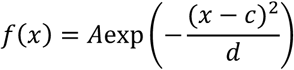

where 𝐴𝐴 is the maximum expression amplitude, 𝑐𝑐 is the age of peak expression, and 𝑑𝑑 is a scale parameter controlling the width of the expression profile. This function approximates biological scenarios where gene activity is highest at a specific developmental stage.

We then assume that part of the genes with age-dependent expression are disease susceptibility genes. Each gene has at most 3 independent (LD *r*^2^<0.01) disease susceptibility loci (sampling without replacement). These loci (*m* in total) collectively explain ℎ^2^ liability to a disease. An individual’s liability for a disease, *L*, is a linear combination of the *m* loci weighted by *w*, plus an additional error term 𝜖𝜖:

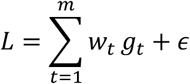

where 𝑔𝑔_𝑖𝑖_ ∈ {0,1,2} is the encoded genotype and the term 𝜖𝜖 represents environmental and unmodeled genetic factors, following a normal distribution 𝜖𝜖 ∼ 𝑁𝑁(0, 𝜎𝜎^2^). Its variance, 𝜎𝜎^2^, is set to ensure that the total genetic component explains a target heritability of liability (ℎ^2^), such that 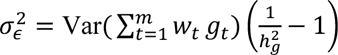. For simplicity, all the weights 𝑤𝑤_𝑡𝑡_ were set to 1.

Disease status was determined using the liability-threshold model^48^. Given a population prevalence 𝐾, individuals with a standardized liability 𝐿𝐿_𝑥𝑥_ exceeding the corresponding threshold 𝑧𝑧 (where 𝑧𝑧 = Φ^−1^(1 − 𝐾𝐾) and Φ^−1^ is the inverse normal cumulative distribution function) were classified as cases.

#### Simulation design and parameterization

Simulations were based on the following parameters:

- Simulated Gene Set: We simulated 20,000 genes. Of these, 500 were modeled with age-dependent expression, while the remaining 19,500 had constant mean expression.
- Disease Susceptibility Genes: A subset of 250 of the 500 age-dependent genes was designated as causal for the simulated phenotype.
- Age-Dependent Expression Parameters: For binary trait simulations, the peak expression age (c) was 20, and the peak width (d) was also 20. For continuous trait simulations, 𝑐𝑐 and 𝑑𝑑 were set to 60 and 20, respectively. Maximum amplitude (𝐴𝐴) was 10 in all cases.
- Genotype Data: We utilized real genotypes for common variants (MAF > 0.05) from 490,541 participants in the UK Biobank.
- eQTL Architecture: Each of the 250 susceptibility genes was regulated by 1 to 5 eQTLs, which collectively explained 25% of its expression variance (ℎ^2^ = 0.25).
- Genetic Contribution to Liability: The 250 susceptibility genes explained 25% of the variance in disease liability (ℎ^2^ = 0.25).
- Disease Prevalence: For binary trait simulations, the population prevalence (𝐾𝐾) was set to 1%.
- Study Cohorts: We simulated a gene expression reference panel of 1,000 individuals across 100 age points (0- 100). For association analysis of binary traits, we drew a separate case-control GWAS cohort of 3,000 cases and 3,000 controls. Association summary statistics were generated using a logistic regression analysis via the KGGA software (https://pmglab.top/kgga). For the association analysis of liability as a continuous trait, we drew a GWAS cohort of 6000 individuals from the simulated population to examine the genetic association by linear regression via KGGA.

#### Evaluation of Type I Error Rate and Statistical Power

We evaluated the performance of tDESE by generating 100 independent GWAS cohorts for each simulation scenario. To assess the type I error rate, we simulated data under the null hypothesis where susceptibility genes had no age-specific expression (i.e., 𝑓𝑓_𝑡𝑡_ (𝑥𝑥) was set to a constant 𝐴𝐴_𝑡𝑡_). The empirical type I error was calculated as the proportion of the 100 simulations that yielded a P-value < 0.05. To assess statistical power, we simulated data under the alternative hypothesis, where the 250 susceptibility genes followed the age-dependent expression patterns described above. Power was calculated as the proportion of the 100 simulations in which tDESE detected a significant association (*P* < 0.05) at the designated peak age.

#### GWAS summary statistics

We collected GWAS summary statistics from public databases for five well-studied neuropsychiatric disorders with distinct age-of-onset patterns: attention deficit hyperactivity disorder (ADHD), schizophrenia (SCZ), major depressive disorder (MDD), bipolar disorder (BIP), and Alzheimer’s disease (AD). We also included four psychological or behavioral traits: neuroticism (NEU), smoking (SMK), sleep duration (SD), and intelligence (IQ). These disorders and traits have large sample sizes, with an average of 442K individuals, ensuring sufficient statistical power for the analyses (see Supplementary Table 1 for details). In all analyses, the major histocompatibility complex (MHC) region was excluded to avoid confounding from its complex linkage disequilibrium structure.

#### Developmental dynamic gene expression profiles

RNA-seq datasets from LIBD_szControl and CommonMind (CMC) were obtained from the PsychENCODE project^19^ (http://www.psychencode.org/). Only healthy postnatal individuals with samples from the dorsolateral prefrontal cortex were included in the current study. Only samples with an RIN greater than 6 were retained to ensure RNA integrity. To reduce the impact of sparse sampling at the extremes of the age distribution, which can cause boundary bias and unstable estimates in the GAM, we applied an age-based trimming procedure before analysis.

Specifically, samples were first sorted by age, and a fixed width sliding window (default width = 5 years) was applied from each end of the age axis to identify the first interval meeting the minimum sample size criterion (default ≥5 samples). The boundaries of these intervals were then used as the start and end points for retention, and only samples within this age range were included in subsequent analyses. This step ensured adequate local sample density at the youngest and oldest ages, thereby stabilizing the local smoothing process in GAM, reducing variance in tail estimates, and improving the reliability of trend and derivative estimates.

In the actual data, the LIBD_szControl dataset originally spanned ages 0–85 years (N=259), which, after quality control, ranged 0–71 years (N=253). The CMC dataset originally spanned ages 17–90 years (N=284), with no change in range or sample size after quality control. For each dataset, genes were retained for downstream analysis if they had non-zero expression in at least five samples, based on the raw count data.

#### Age at diagnosis from epidemiological surveys

For four psychiatric disorders, namely attention deficit hyperactivity disorder, schizophrenia, major depressive disorder and bipolar disorder, we referred to a large-scale meta-analysis of epidemiological studies on mental disorders^9^ to obtain the interquartile range (Q1–Q3) of the age-at-onset distribution, which was taken as the peak age range for each disorder. For Alzheimer’s disease, as the vast majority of cases occur after the age of 65 years^18^, we adopted the conventional definition of 65–100 years as its high-risk age range.

#### Identification of time-sensitive genes at high-risk disease ages and functional enrichment analysis

To identify genes specifically highly expressed during disease high-risk age periods, we calculated the Spearman’s rank correlation coefficient between each gene’s expression levels across different ages and the disease enrichment scores at corresponding ages. A higher correlation coefficient indicates stronger age-specific expression of the gene during the disease high-risk period. Genes with the highest correlation coefficients were defined as “temporally sensitive genes” at high-risk disease ages. In subsequent analyses, the top 200 temporally sensitive genes ranked by correlation coefficient were selected for functional enrichment analysis, including both the general Gene Ontology (GO) database and the SynGO database focused on synaptic function.

GO enrichment analysis was performed using the API provided by g:Profiler (https://biit.cs.ut.ee/gprofiler)^49^, selecting three GO categories: Biological Process (GO:BP), Cellular Component (GO:CC), and Molecular Function (GO:MF). Enrichment significance was assessed by Fisher’s exact test, and multiple testing correction was applied using the default g:SCS algorithm. SynGO enrichment analysis was conducted via the SynGO online platform^25^ (https://www.syngoportal.org/), using Fisher’s exact test for significance and false discovery rate (FDR) correction for multiple testing.

#### Sex-specific analysis of disease-associated ages

We stratified the developmental dynamic expression profiles by sex and performed tDESE analysis separately to identify sex-specific high-risk ages for disease. Due to insufficient female samples under the age of 35 in the CMC dataset after sex stratification, these samples did not pass quality control (see Supplementary Figure 1), and therefore, this dataset was not used for sex-stratified tDESE analysis. In the LIBD_szControl dataset, the reduced sample size after sex stratification led to a loss of statistically significant enrichment signals for some diseases.

#### Similarity analysis of high-risk age patterns across phenotypes

To assess the similarity of high-risk age patterns among different phenotypes, we calculated the Spearman’s rank correlation coefficient between enrichment scores of each phenotype across different ages. A higher correlation coefficient indicates a stronger concordance between two phenotypes in their high-risk age patterns.

#### Gene prioritization and evaluation based on age-specific expression

To evaluate whether prioritizing disease risk genes based on age-specific expression during peak disease ages improves gene prioritization performance, we ranked disease-associated genes according to their specificity of expression at each age. Disease-associated genes were defined as those with Benjamini-Hochberg adjusted P values below 0.01. If more than 1000 genes met this threshold, only the top 1000 genes by P-value were retained.

Next, we constructed a benchmark disease gene set using the Open Targets Platform (release 25.06)^50^, which is a comprehensive tool supporting the systematic identification of potential therapeutic targets and their prioritization. For our benchmark, we selected genes with an overall association score ≥0.4, reflecting moderate or higher aggregated evidence, as well as genes with drug evidence recorded in ChEMBL (ChEMBL score >0). The union of these two sets was used as a high-confidence reference set for disease genes. Based on this benchmark, we performed receiver operating characteristic (ROC) analysis and calculated the area under the curve (AUC) for gene prioritization at each age. This evaluation assessed the accuracy of gene prioritization at different ages and identified the age at which prioritization performance was optimal.

## Data availability

GWAS summary statistics for the five neuropsychiatric disorders were obtained from the Psychiatric Genomics Consortium (PGC, https://pgc.unc.edu/for-researchers/download-results/), and those for four psychological and behavioral traits were obtained from the GWAS Catalog (https://www.ebi.ac.uk/gwas/). Developmental dynamic expression data for the human dorsolateral prefrontal cortex were retrieved from the PsychENCODE project^19^ via NIMH Data Archive (NDA). Genotype data for the European population from Phase 3 of the 1000 Genomes Project were downloaded from ftp://ftp-trace.ncbi.nih.gov/1000genomes/ftp/release/20130502. Benchmark disease genes were obtained from the Open Targets Platform (https://platform.opentargets.org/). In the simulation study, the common variants (MAF>0.04) in whole genome sequencing datasets of the UK Biobank (https://www.ukbiobank.ac.uk)^12^ were accessed through a collaboration under application ID 86920. Genotype data in VCF.GZ format were obtained under Field ID 24310. These data are available to bona fide researchers upon application to the UK Biobank.

## Code availability

The functionality of tDESE has been implemented within the Java-based KGGSUM analysis platform (https://pmglab.top/kggsum). Source code for the analyses and visualizations presented in this study is available at https://github.com/chaoxue-gwas/tDESE.

## Acknowledgements

This work was supported by the National Natural Science Foundation of China (grant numbers 32470683, 32470680, 32300500, and 32170637). We thank NDA for approving our data access for PsychENCODE project (DAR ID 23214). NDA is a collaborative informatics system created by the National Institutes of Health to provide a national resource to support and accelerate research in mental health. This manuscript reflects the views of the authors and may not reflect the opinions or views of the NIH or of the Submitters submitting original data to NDA.

## Author contributions

C.X., X.Y., and M.L. conceived the study. C.X., X.Y., L.Y., and B.T. developed the statistical methods. L.J. performed the simulation analyses. M.L. and H.G. applied and provided the dataset for analysis. C.X., X.Y. and H.G. conducted data processing, computational analyses, and biological interpretation of the results. C.X., Y.L., L.Z. and M.L. developed the software. C.X., X.Y., L.J., and M.L. wrote the manuscript. C.X. and M.L. secured funding. M.L. provided supervision throughout the study.

## Supplementary information

Supplementary Information: Supplementary Figure 1-2 and Table 1.

Supplementary Data: Supplementary data tables.

