## Supplementary Figure 1-2 and Table 1 for "Developmental expression of risk genes implicates the age of onset for neuropsychiatric disorders"

Supplementary Figures

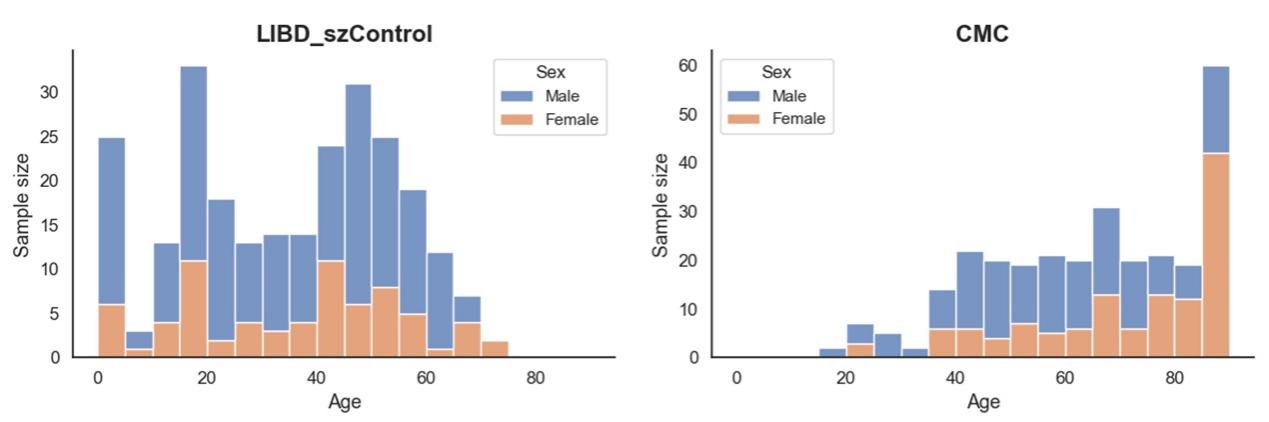

**Supplementary Figure 1. Sex and age distribution of samples in the developmental dynamic expression dataset.** Age is shown on the x-axis, starting from 0 years and grouped into 5-year intervals, while the y-axis indicates the number of samples. Colors represent sex.

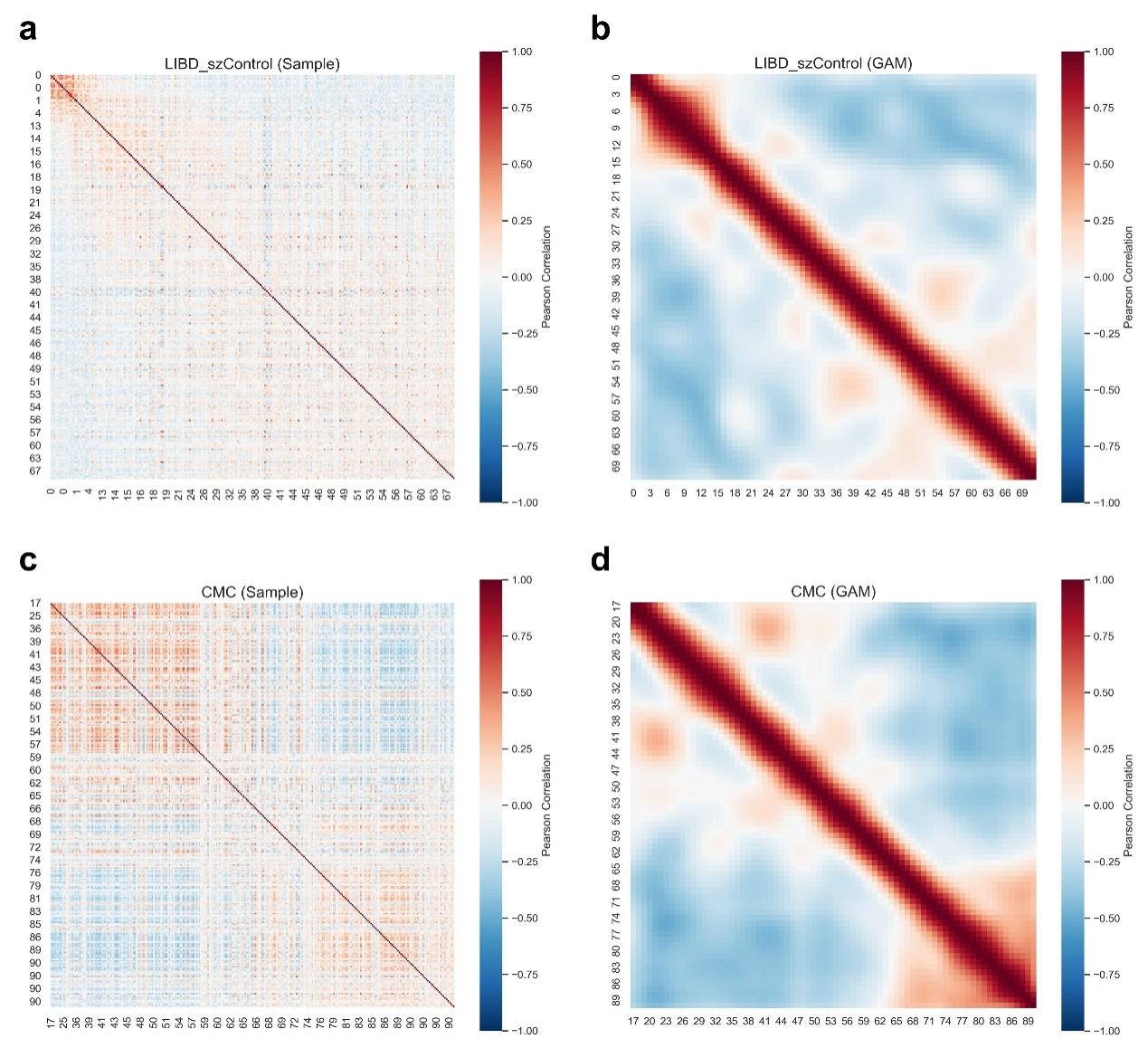

**Supplementary Figure 2. Heatmaps of Pearson correlation coefficients based on either age-stratified samples or GAM-predicted expression.** (a, c) Correlation matrices calculated from the original expression profiles of the samples, with samples ordered by increasing age. (b, d) Correlation matrices calculated from GAM-predicted expression profiles, with samples similarly ordered by age.

Supplementary Tables

Note: Only the title and legend are shown here; the complete dataset can be found in the corresponding table of the supplementary Excel file.

**Supplementary Table 1. GWAS summary statistics of nine complex diseases and traits.**
